# Litterbox - A gnotobiotic zeolite-clay system to investigate Arabidopsis-microbe interactions

**DOI:** 10.1101/2020.01.28.922625

**Authors:** Moritz Miebach, Rudolf Schlechter, John Clemens, Paula E. Jameson, Mitja N.P. Remus-Emsermann

**Affiliations:** School of Biological Sciences, University of Canterbury, Christchurch, New Zealand; Biomolecular Interaction Centre, University of Canterbury, Christchurch, New Zealand

## Abstract

Plants are colonised by millions of microorganisms representing thousands of species with varying effects on plant growth and health. The microbial communities found on plants are compositionally consistent and their overall positive effect on the plant is well known. However, the effects of individual microbiota members on plant hosts and *vice versa*, as well as the underlying mechanisms remain largely unknown. Here, we describe ‘Litterbox’, a highly controlled system to investigate plant-microbe interactions. Plants were grown gnotobiotically on zeolite-clay, an excellent soil replacement that retains enough moisture to avoid subsequent watering. Plants grown on zeolite phenotypically resemble plants grown under environmental conditions. Further, bacterial densities on leaves in the Litterbox system resembled those in temperate environments. A PDMS sheet was used to cover the zeolite, thereby significantly lowering the bacterial load in the zeolite and rhizosphere. This reduced the likelihood of potential systemic responses in leaves induced by microbial rhizosphere colonisation. We present results of example experiments studying the transcriptional responses of leaves to defined microbiota members and the spatial distribution of bacteria on leaves. We anticipate that this versatile and affordable plant growth system will promote microbiota research and help in elucidating plant-microbe interactions and their underlying mechanisms.

## Introduction

Plants offer three different habitats to microbes: the endosphere, the rhizosphere, and the phyllosphere. The endosphere encompasses the habitat formed by internal tissues of plants, whereas the rhizosphere and phyllosphere encompass the surfaces of belowground and aboveground plant organs, respectively. Plants host remarkably diverse and complex, yet structured, microbial communities, collectively referred to as the plant microbiota [1–3]. Due to the microbiota’s diversity and complexity, it is not surprising that the traditional view of host-microbe interactions focussing on plant pathogens, nitrogen-fixing rhizobacteria, and phosphate-mobilizing mycorrhizal fungi, has recently shifted to a holistic view considering the plant and its associated microbiota as a metaorganism or holobiont [4–6]. It is widely recognised that members of the microbiota assist in nutrient uptake, promote growth, and protect against biotic and abiotic stresses [7–14]. To harness these positive impacts of plant-associated microbiota, the use of synthetic microbiota has been proposed [15–17]. The prospect of using synthetic microbial communities to promote sustainable agriculture is leading to a growing appreciation of plant microbiota research.

Generally, plant microbiota research aims to understand 1) plant-microbe and microbe-microbe interactions ranging from the individual microorganism to the microbial community resolution, 2) the underlying molecular mechanisms of these interactions, and 3) their contribution to microbial community structure. Thus far, most of our understanding of microbiome structure and function is derived from observations at the whole community level (16S rDNA and ITS sequencing, meta-genomics, -transcriptomics, and -proteomics, etc.) [18–22]. Such observations showed that microbial community structure is governed by many factors, including abiotic factors, physicochemical properties of the leaf, plant-microbe and microbe-microbe interactions [2]. However, owing to the multitude of factors affecting microbial community structure, their effects are often convoluted. Therefore, reductionist approaches are needed to disentangle the complexity and to allow a shift from association to causation [23]. Bai et al. (2015) have built an extensive indigenous bacterial culture collection covering phyllosphere and rhizosphere inhabitants of *Arabidopsis thaliana*, enabling the research community to study the functions of individual microbiota members in synthetic communities [24]. While synthetic communities might not perfectly mimic nature, they allow for testing of causality either by bottom-up (adding microorganisms to low complexity synthetic communities) or drop-out (excluding single strains from rather complex synthetic communities) experiments [23,25,26]. Furthermore, to fully understand plant-microbe and microbe-microbe interactions in the heterogeneous phyllosphere, the interactions need to be studied at micrometer resolution, the scale relevant to microbes [1].

A deeper understanding of the driving factors of microbial community structure will enable us to engineer plant protective synthetic microbial communities with increased resilience. In addition, to engineer plant protective synthetic microbial communities that are superior to individual biocontrol strains, knowledge about the varying modes of plant protection by biocontrol strains and the compatibility of these biocontrol strains is essential. For example, it was shown that *Pseudomonas fluorescens* A506 can alleviate the biocontrol activity of *Pantoea vagans* C9-1, whereas each is individually effective as a biocontrol strain against *Erwinia amylovora*, the causal agent of fire blight [27]. *Pseudomonas fluorescens* A506 produces an extracellular protease that degrades the peptide antibiotics produced by *Pantoea vagans* C9-1 [28]. Therefore, to fully harness the power of the microbiome, our understanding of the varying modes of biocontrol activity needs to be extended by studying the underlying mechanisms of plant-microbe and microbe-microbe interactions. To do so, hypothesis-driven research needs to be conducted in a highly controlled, yet adjustable gnotobiotic, i.e. otherwise sterile, system.

Highly controlled and reproducible growth conditions are needed as abiotic factors can influence the plant’s response to its microbial colonisers [29,30]. Furthermore, pH is a strong driving factor of soil microbial communities [31,32]. Controlling abiotic factors alleviates confounding effects of these on microbial community structure, and will thus allow for time-course experiments, which are important as plant responses to microbial colonisers are dynamic [33,34]. The order in which microbes are introduced, as well as genetic alterations of the host or the microbes allows for a mechanistic understanding of plant-microbe and microbe-microbe interactions.

Arabidopsis can be grown gnotobiotically under tissue culture conditions on a variety of different substrates. Phytoagar is widely used, allows for reproducible growth, and is easy to handle. Phytoagar-based systems have also been successfully used for plant-microbe interaction studies [9,35]. However, aeration of the root system is sub-optimal and nutrients are non-uniformly delivered to the root surface, rendering the system highly artificial [36]. Hydroponics and aeroponics allow for a uniform distribution of nutrients and O_2_, but the lack of structure leads to abnormal root phenotypes [36]. However, this can be circumvented by the use of filter paper in a hydroponic system [37]. Nevertheless, these systems are often labour intensive to set up. Furthermore, plants grown in such systems are exposed to high to very-high humidity, which is not feasible to modulate over a wide range. The microclimate in which a plant grows severely affects cuticle composition leading to differences in water permeability and leaf wettability and finally to shifts in microbial community structure [38–40]. Furthermore, humidity has been described to be one of the main drivers for bacterial colonisation on plants [41]. With the aim to minimise experimental artifacts and mimic natural conditions, soil has been used in axenic plant growth systems. However, sterilisation of soil is difficult and usually accompanied by structural and geochemical changes [42,43]. Recently, calcined clay was used as an alternative to soil, rendering a similar texture to soil, but adjustable in its nutrient profile [44,45]. However, the desorptive properties of calcined clay can lead to toxic levels of labile Mn ions [46].

Here we introduce ‘Litterbox’ a gnotobiotic plant growth system that uses zeolite as a soil-like substrate that mimics environmental conditions. Zeolite is easy to sterilise, its porous structure ensures root aeration, it has an excellent water- and nutrient-holding capacity, it is known for its slow-release of nutrients, including Mn, thereby alleviating the risk of Mn toxicity [47,48], and has previously been used in other plant growth systems, even on space missions [49]). Additionally, by adding a PDMS sheet as a protective layer onto the zeolite, cross-inoculation of phyllosphere and rhizosphere can be significantly reduced. Zeolite can be easily and affordably sourced from commercial “cat litter”.

## Materials and Methods

### Plant growth

Seeds of *Arabidopsis thaliana* (Col-0) were surface sterilised according to Lundberg et al. (2012) [50]. Depending on the experiment, seeds were either sown directly on zeolite (sourced from cat litter – Vitapet, Purrfit Clay Litter), on ½ Murashige and Skoog medium (MS, including vitamins, Duchefa) 1% phyto agar (Duchefa) plates, pH 5.9, or on pipette tips (10 µl or 200 µl) filled with ½ MS 1% phyto agar, pH 5.9. Seeds were germinated in a CMP6010 growth cabinet (Conviron) with 11 h of light (150 - 200 µmol m^−2^ s^−1^) and 85% relative humidity. Seedlings germinated on agar plates or agar-filled pipette tips were transferred seven days after sowing to autoclaved plant tissue culture boxes (Magenta vessel GA-7) filled with either 90 g fine ground zeolite or 70 g coarse zeolite covered by 20 g finely ground zeolite, soaked with 60 ml of ½ MS including vitamins, pH 5.9. Fine ground zeolite was obtained using a jaw crusher (Rocklabs, Boyd Crusher). The zeolite was covered with a polydimethylsiloxane (PDMS) sheet (thickness 0.5 – 1.5 mm) to reduce inoculation of the growth substrate. Lids of the plant tissue culture boxes contained four holes for gas exchange (9 mm diameter), which were covered by two pieces of micropore tape (3M) to ensure axenic conditions. Plants were grown in a growth room at 21 °C and 11 h of light (120 – 200 µmol m^−2^ s^−1^). Plant tissue culture boxes were put into transparent plastic containers (Sistema) closed with cling wrap to maintain high relative humidity. After inoculation, the plant tissue culture boxes were kept in a CMP6010 growth cabinet with 11 h of light (150 - 200 µmol m^−2^ s^−1^) and 85% relative humidity. High batch to batch variation in readily available ions was reported in calcined clays leading to heavy metal toxicity [46]. We also observed plant phenotypes resembling heavy metal toxicity in one out of ten batches of cat litter. It is thus advisable to test the growth of a few plants when switching to a new batch of cat litter.

### Fabrication of PDMS sheets

4 g of catalyst was added to 40 g of PDMS and the solution vigorously mixed using a plastic rod. To release trapped air bubbles, the mix was vacuumed, and the vacuum was released several times until no more air bubbles appeared. Two A4 acetate sheets (OfficeMax, Overhead Projector Transparency Film), were placed on a flat surfaced sandwich press (Sunbeam, GR8450B) with an area of approximately 720 cm^2^. The PDMS was poured on top and evenly distributed with a spatula. The top of the sandwich press was lowered until sitting approximately 9 mm above the PDMS without touching. The PDMS was cured at 60 °C for 1 h. Then the PDMS was carefully pulled from the acetate sheets and cut to 6 × 6 cm squares. A hole punch was used to punch four holes into each PDMS sheet, to allow the growth of four plants per plant tissue culture box.

### Plant inoculation

*Sphingomonas melonis* FR1 and *Pseudomonas syringae* DC3000 (Table 1) were cultivated at 30 °C on R2A (HIMEDIA) or King’s B (HIMEDIA) media plates, respectively. In case of subsequent plant gene expression analysis the bacterial strains were cultivated on minimal media [51] agar plates containing 0.1% pyruvate as a carbon source to prevent the contamination of plants with growth media-derived cytokinins that may impact plant responses [52]. Bacterial suspensions were prepared from bacterial colonies suspended in phosphate buffered saline (PBS, 0.2 g l^−1^ NaCl, 1.44 g l^−1^ Na_2_HPO_4_ and 0.24 g l^−1^ KH_2_PO_4_) and washed twice via centrifugation at 4000 x *g* for 5 min followed by discarding the supernatant and adding PBS. The bacterial suspensions were then diluted to a desired optical density (OD = 0.025). 200 µl of bacterial solution was sprayed per plant tissue culture box using an airbrush spray gun (Pro Dual Action 3, 0.2 mm nozzle diameter). To obtain a homogeneous coverage, the distance between the airbrush spray gun and the plants was increased by stacking a bottomless plant tissue culture box onto the plant tissue culture box, containing the plants being spray-inoculated.

**Table 1:** List of bacterial strains used in this study.

| Bacterial strain | Reference | Fluorescence |
| --- | --- | --- |
| *Sphingomonas melonis* Fr1 | [55] | n.a. |
| *Pseudomonas syringae* DC3000 | [56] | n.a. |
| *Sphingomonas melonis* Fr1 | This work | mTurquoise2 (cyan) |
| *Pseudomonas syringae* DC3000 | This work | mScarlet-I (red) |

To co-inoculate *S. melonis* Fr1::mTurquoise2 and *P. syringae* DC3000::mScarlet-I (Table 1) onto *A. thaliana* leaves, a bacterial suspension was prepared as described above to a final optical density of 0.005 (1:1 OD ratio) before being airbrushed onto 47 days-old Arabidopsis. Fluorescently-tagged strains were developed as described previously [53,54].

### Enumeration of bacteria recovered from varying environments

Bacterial cell numbers in different environments (phyllosphere, rhizosphere, zeolite) were determined. Aboveground plant parts were carefully detached from the belowground parts and placed individually in 1.5 ml tubes. To sample the rhizosphere, the zeolite containing the roots was poured out and the roots were cut and placed individually in 1.5 ml tubes. A spatula was then used to collect between 100 and 300 mg of zeolite, placed individually in 1.5 ml tubes. After determining plant or zeolite sample weight, 1 ml PBS with 0.02% Silwet L-77 (Helena chemical company) was added to each tube. The bacteria were then dislodged from the sample via shaking twice for 5 min at 2.6 m/s in a tissue lyser (Omni Bead Ruptor 24), followed by sonication in a water bath for 5 min. 3 µl spots of tenfold dilutions of each sample were spotted onto R2A plates. Colony forming units (cfu) were counted after incubation of the plates at 30 °C. In the case of co-inoculation experiments, *S. melonis* Fr1::mTurquoise2 and *P. syringae* DC3000::mScarlet-I were selected on R2A plates containing tetracycline (15 µg/ml) or gentamicin (20 µg/ml), respectively.

### Gene expression analysis

Six-week-old plants were spray-inoculated with either *P. syringae* DC3000, *S. melonis* Fr1 or PBS (mock control), and harvested over time (3 h, 9 h, 48 h). Four mature leaves per plant were collected in an RNase-free 1.5 ml tube and then immediately frozen in liquid N_2_. Six plants were sampled per treatment and time point. Flash frozen samples were ground using Teflon pestles in the collection tube. RNA extraction was performed using the Isolate II RNA Plant kit (Bioline). During RNA extraction two samples were combined by using the same lysis buffer on two samples to make up one biological replicate. Purity and quantity of the RNA sample was determined using the Nanodrop (ND-1000 Spectrophotometer). Integrity of the RNA sample was assessed via gel electrophoresis. Approximately 1 µg of RNA was used for cDNA synthesis and for the no Reverse Transcriptase (noRT) control, using the VitaScript First strand cDNA synthesis kit (Procomcure Biotech). RT-qPCR was performed using the 2x ProPlant SYBR Mix (Procomcure Biotech) in 15 µl reaction volumes with 0.2 µM of each primer and 0.001 g/l of initial RNA in the cDNA mix. QPCRs were run using the recommended protocol for 2x ProPlant SYBR Mix (Procomcure Biotech) on a Rotor-Gene Q (Qiagen). Technical triplicates were performed for each biological replicate. The ROX dye, present in the 2x ProPlant SYBR Mix (Procomcure Biotech), was used to normalise for master mix variation between tubes. A mix of equal amounts of all cDNAs was used for normalisation between runs. mRNA concentrations were calculated using Equation 1.

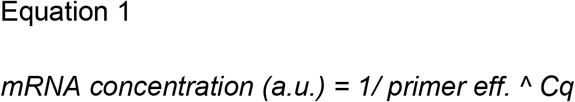

Primers that were first used in this study were designed using either the ‘Universal ProbeLibrary Assay Design Center’ (Roche) or ‘primer-blast’ (NCBI). Primer efficiencies were determined via serial template dilutions [57]. The mRNA concentration of each target gene was then normalised against the mean mRNA concentration of five stably expressed, previously described reference genes (Table 2, [58,59]). Next, the normalised mRNA concentration of each treatment (*P. syringae* DC3000 and *S. melonis* Fr1 inoculation) was normalised against the mean normalised mRNA concentration of mock treated samples, to emphasise treatment related changes in gene expression.

**Table 2:**
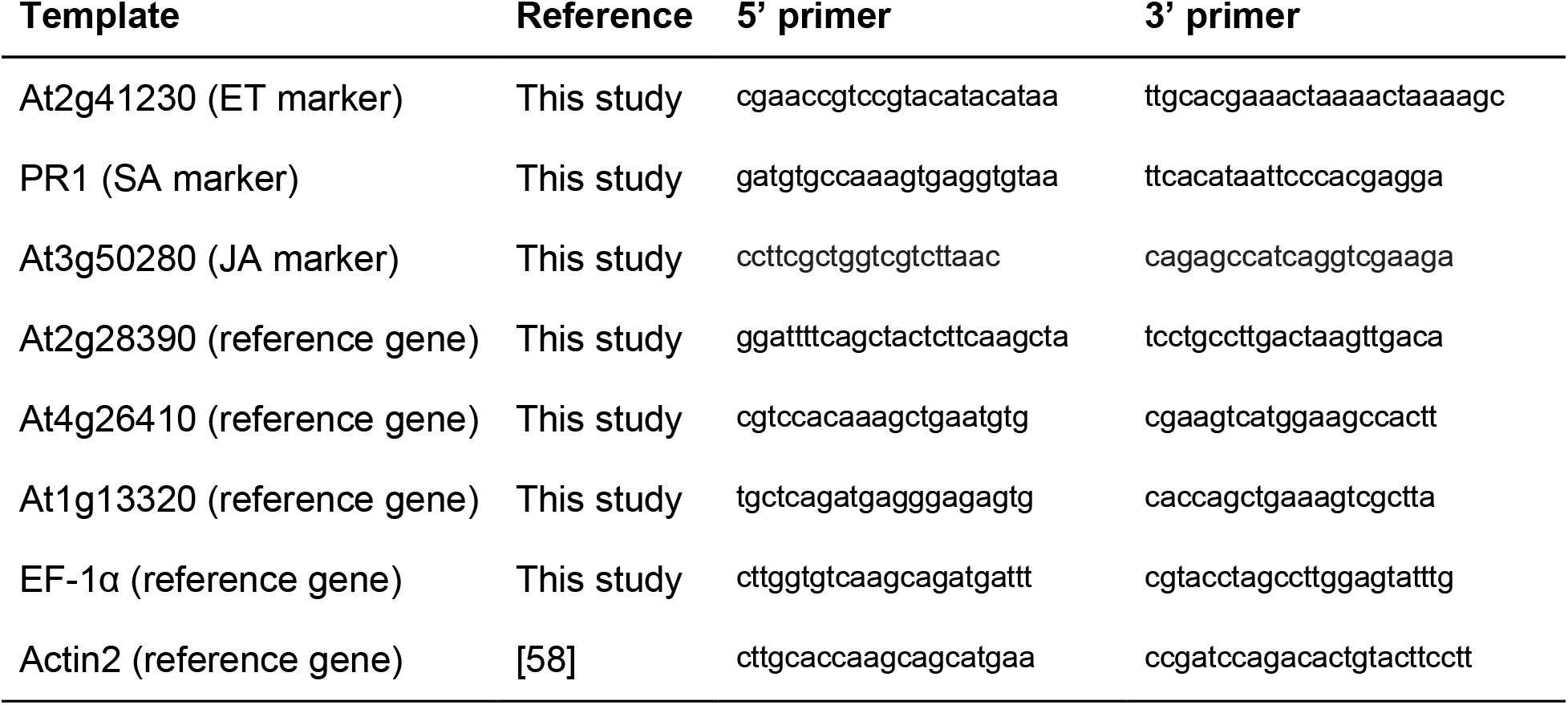
List of primers used in this study.

### Microscopy and image processing

Bacteria were recovered 14 days post-inoculation from the abaxial side of co-inoculated Arabidopsis leaves using the cuticle tape lift procedure described previously [60]. The procedure involves placing one side of the leaf onto double sided adhesive tape and carefully stripping them off again. Cuticle tape lifts allow the recovery of phyllosphere bacteria from leaves without redistributing their spatial location.

Microscopy was performed using the Zeiss AxioImager.M1 fluorescent widefield microscope at 100x magnification (EC Plan-Neofluar 100x/1.30 PH3 Oil M27 objective) equipped with Zeiss filter sets 38HE (BP 470/40-FT 495-BP 525/50) and 43HE (BP 550/25-FT 570-BP 605/70) for the detection of *S. melonis* Fr1::mTurquoise2 and *P. syringae* DC3000::mScarlet-I, respectively. Acquisition of 3D images was achieved using an Axiocam 506 and the software Zeiss Zen 2.3. FIJI/ImageJ was used for image processing [61]. To improve the signal-to-noise ratio and the depth of field, background subtraction was used before images were z-stacked using maximum intensity projection. The contrast of each resulting image was enhanced.

## Results and discussion

Arabidopsis plants were grown in tissue-culture boxes on either agar or zeolite, to compare the suitability of the substrates for gnotobiotic plant growth. To that end, seven-day-old seedlings, that had been germinated on cut, agar-filled pipette tips were transferred to the respective substrate and plant growth was evaluated over a period of three weeks. Both substrates were covered by a PDMS sheet to reduce the cross-inoculation of phyllosphere and rhizosphere bacteria (Figure 1A,B). Plant growth in the Litterbox system was optimised beforehand to determine optimal amounts of zeolite per tissue-culture box, optimal zeolite granularity and optimal buffering of growth media (Figure S1). Growth of Arabidopsis was markedly different on agar compared with zeolite. The most prominent differences were observed in the rootzone. Tight root networks were observed on the bottoms (Figure 1E) and the sides (Figure 1I) of the Litterbox. In contrast, in the agar system only a few roots penetrated into the agar and those that penetrated did not grow to the bottom of the tissue-culture box (Figure 1F,J). Most of the roots grew on top of the agar (Figure 1H), which is unnatural and did not occur in the Litterbox system (Figure 1G). Furthermore, roots on the agar were noticeably curled (Figure 1H), which is likely linked to a continuous root growth reorientation by the environment-sensing root cap [62]. Predominant growth on top of the agar might be linked to a humidity gradient with the highest water potential on top of the agar, potentially caused by the PDMS sheet [63]. In general, humidity can be adjusted in the Litterbox system by changing the ratio of media to zeolite, whereas in the agar system humidity cannot be adjusted without the agar drying out. Moreover, in agar-based systems the bacterial load in the phyllosphere is usually around 10^8^ cfu/g [9,64,65], which is one to two magnitudes greater than under temperate environmental conditions which have been reported to be 10^6^-10^7^ cfu/g [66,67].

**Figure 1:**
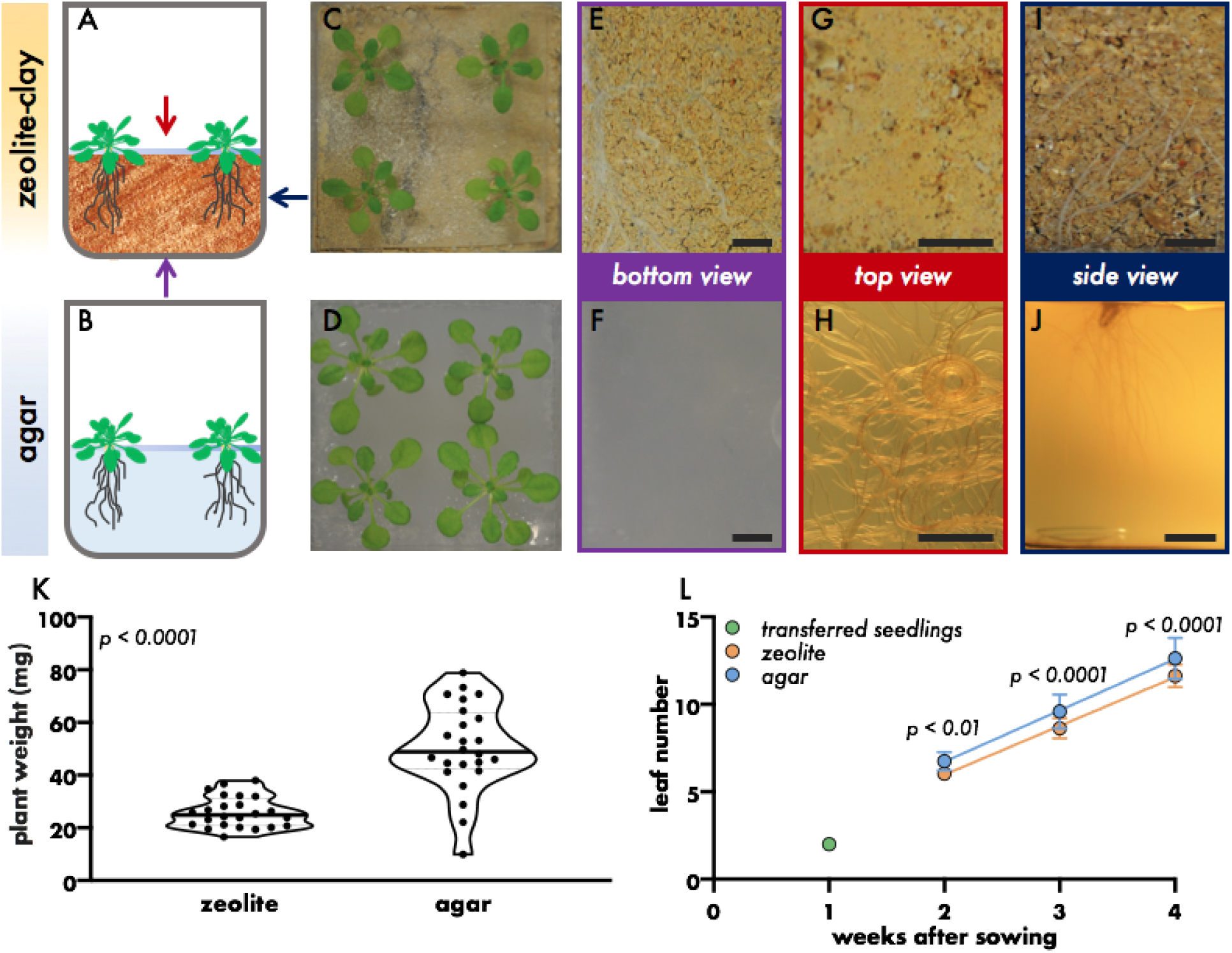
The effect of growth substrate on plant phenotype. **A, B.** Illustration of general growth setup. Blue bar represents PDMS sheet. Colored arrows indicate the point of view onto the plant growth box. **C, D.** Representative images of four weeks old plants grown on zeolite (**C**) or agar (**D**). **E-J.** Representative images of roots of six week old plants grown on zeolite (**E,G,I**) or agar (**F,H,J**). View on growth box bottom (purple arrow, **E,F**), top of growth medium (red arrow, **G,H**) and side of growth box (dark blue arrow, **I,J**), bar scale = 5 mm. **K.** Plant weight (fresh weight aboveground plant parts) of four-week-old plants grown on zeolite or agar. Dots represent individual samples, thick bar represents median, dotted bars represent quartiles, Mann-Whitney test. **L.** Number of leaves per plant at different weeks after sowing of plants grown either on zeolite or agar. Seedlings were transferred one week after sowing to either zeolite or agar. Filled circles represent mean of 24 plants, error bars represent standard deviation, colored lines represent linear regression lines, Sidak’s multiple comparison test.

Four-week-old plants grown on zeolite were smaller (Figure 1C,D), had significantly fewer leaves (Figure 1L) and were significantly lighter than plants grown on agar (Figure 1K). Noticeably, there was no significant difference in the growth rate of plants grown on agar or zeolite, from one week after transfer to tissue-culture boxes (two weeks after sowing) until harvest (Figure 1L; linear regression, P = 0.343, ANCOVA). Therefore, the difference in plant weight and developmental stage was likely caused by a short halt in growth after seedling transfer to zeolite, due to adaptation to a new environment. More importantly, plants grown on zeolite showed a lower degree of variation than plants grown on agar (Coefficient of variation: zeolite = 22.77%, agar = 32.92%), which makes the Litterbox system advantageous for predictable and low variability growth during experiments, which should result in fewer false positive and negative measurements (Figure 1K).

Additionally, plants that were pre-grown on cut, agar-filled pipette tips (Figure 2A) prior to transfer to zeolite exhibited a lower degree of variation than plants of seeds that were directly sown on zeolite and plants that were transferred without pipette tip (Figure 2B, Coefficient of variation: sown = 34.7%, transferred seedling = 42.28%, transferred seedling on tip = 22.77%). Almost all of the seedlings that were pre-grown in and then transferred with the pipette tip survived the transfer to zeolite, whereas only 61.1% of the seedlings that were transferred from an agar plate without pipette tips survived (Figure 2C). Only 84.7% of seeds sown directly on zeolite germinated. Therefore, it is highly advisable to sow and pre-grow the seeds and seedlings in agar-filled pipette tips, especially when working with transgenic or mutant plant lines that exhibit low germination rates. Different sized pipette tips filled with varying agar strength were tested. Plants pre-grown on pipette tips filled with 1% agar were significantly bigger than plants grown on pipette tips with 0.6% agar strength (Figure S2A). Further, plants pre-grown in 200 µl pipette tips exhibited poor survival rates at 0.6% agar (62.5% survived plants), but not at 1% agar (95.83% survived plants) after transfer to zeolite (Figure S2B).

**Figure 2:**
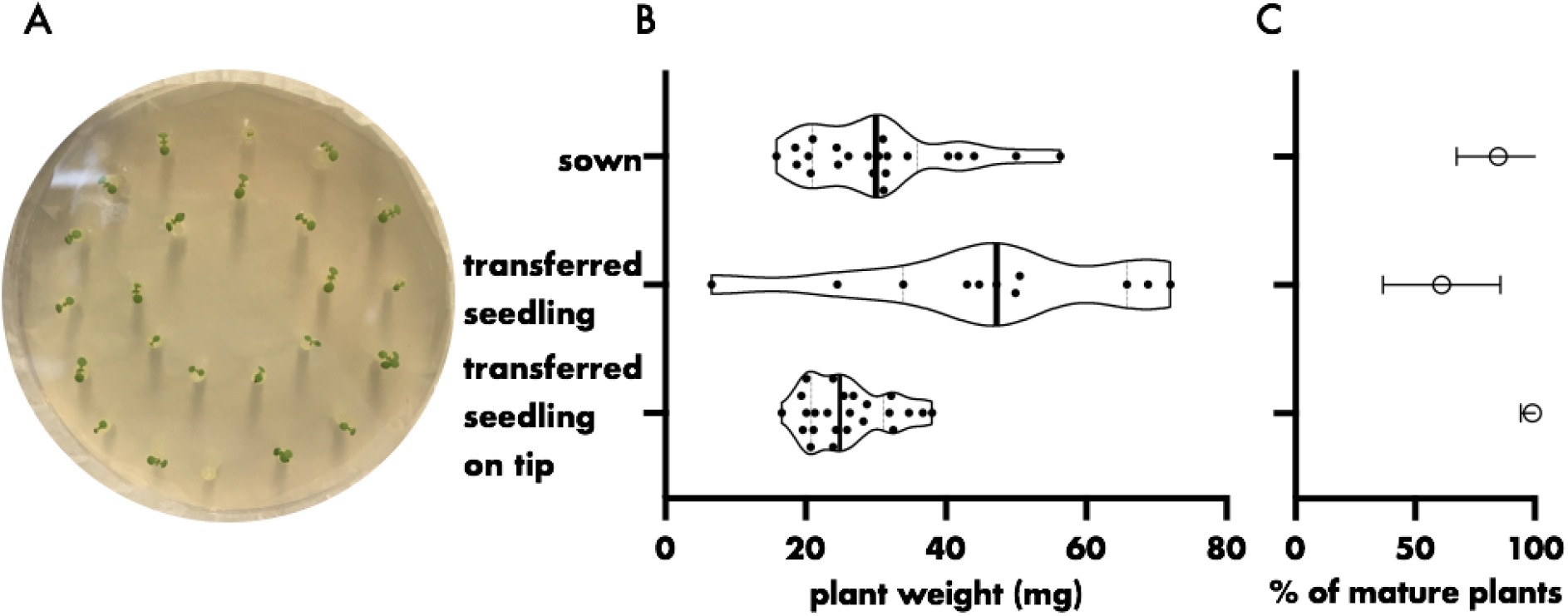
The effect of different sowing strategies on germination rate/seedling survival and plant weight. **A.** Representative image of seedlings on agar-filled pipette tips seven days after sowing. **B,C.** Plant weight (fresh weight aboveground plant parts) (**B**) and the percentage of mature plants per seed/seedling (**C)** after four weeks of growth. Seeds were either directly sown on zeolite, germinated on agar before transfer to zeolite, or germinated on agar-filled pipette tips before transfer to zeolite. Seedlings were transferred seven days after sowing. Hollow dots represent sample mean, error bars depict standard deviation, filled dots mark individual plants, thick bar represents median, dotted bars represent quartiles.

As mentioned above, the bacterial load on leaves in agar-based systems is up to two magnitudes higher than under environmental conditions [9,64–66]. In the Litterbox system, bacteria reproduced in the phyllosphere, their growth over time was tracked and eventually they reached densities of 10^7^ bacteria per g leaf (fresh weight), which is comparable to bacterial densities found on leaves in nature (Figure 3).

**Figure 3:**
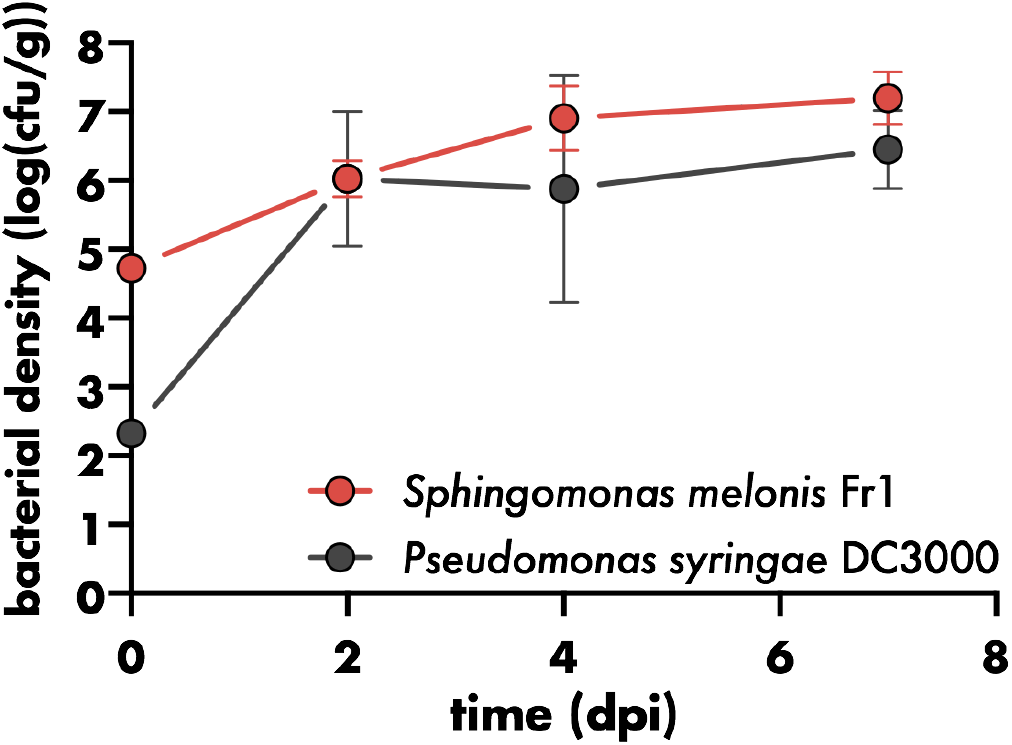
Bacterial growth in the phyllosphere. Bacterial density on aboveground plant parts of six-week-old plants relative to plant weight (fresh weight of aboveground plant parts). Filled circles represent mean of four plants, error bars represent standard deviation.

The Litterbox system can be used for a variety of different study designs. Besides phyllosphere studies, it can also be used for rhizosphere studies, as bacteria reproduced in the rhizosphere and the zeolite (Figure 4). However, other systems such as the recently published FlowPot system, which allows rapid changes in media composition, might be advantageous for studies focussing on the rhizosphere [68]. For phyllosphere studies, the Litterbox system is the optimal choice, not only because Litterbox-grown Arabidopsis plants resemble Arabidopsis plants grown in temperate environments with regard to phenotype and bacterial densities found on leaves, but also because the use of a PDMS sheet as a protective layer reduces cross-inoculation between phyllosphere and rhizosphere. This is especially advantageous for studies that aim to disentangle local from systemic plant responses to microbial colonisation. Plant growth promoting rhizobacteria, for example, were shown to confer broad-spectrum resistance to the entire plant upon rhizosphere colonisation by a process called induced systemic resistance (ISR) [69,70]. Notably, to induce ISR these bacteria need to be present in high densities above 10^5^ bacteria per gram root [71]. The presence of a PDMS sheet significantly reduced the bacterial load in the rhizosphere and zeolite after spray-inoculation of the phyllosphere by an order of magnitude (Figure 4). Plant growth was not affected by the PDMS sheet (Figure S3).

**Figure 4:**
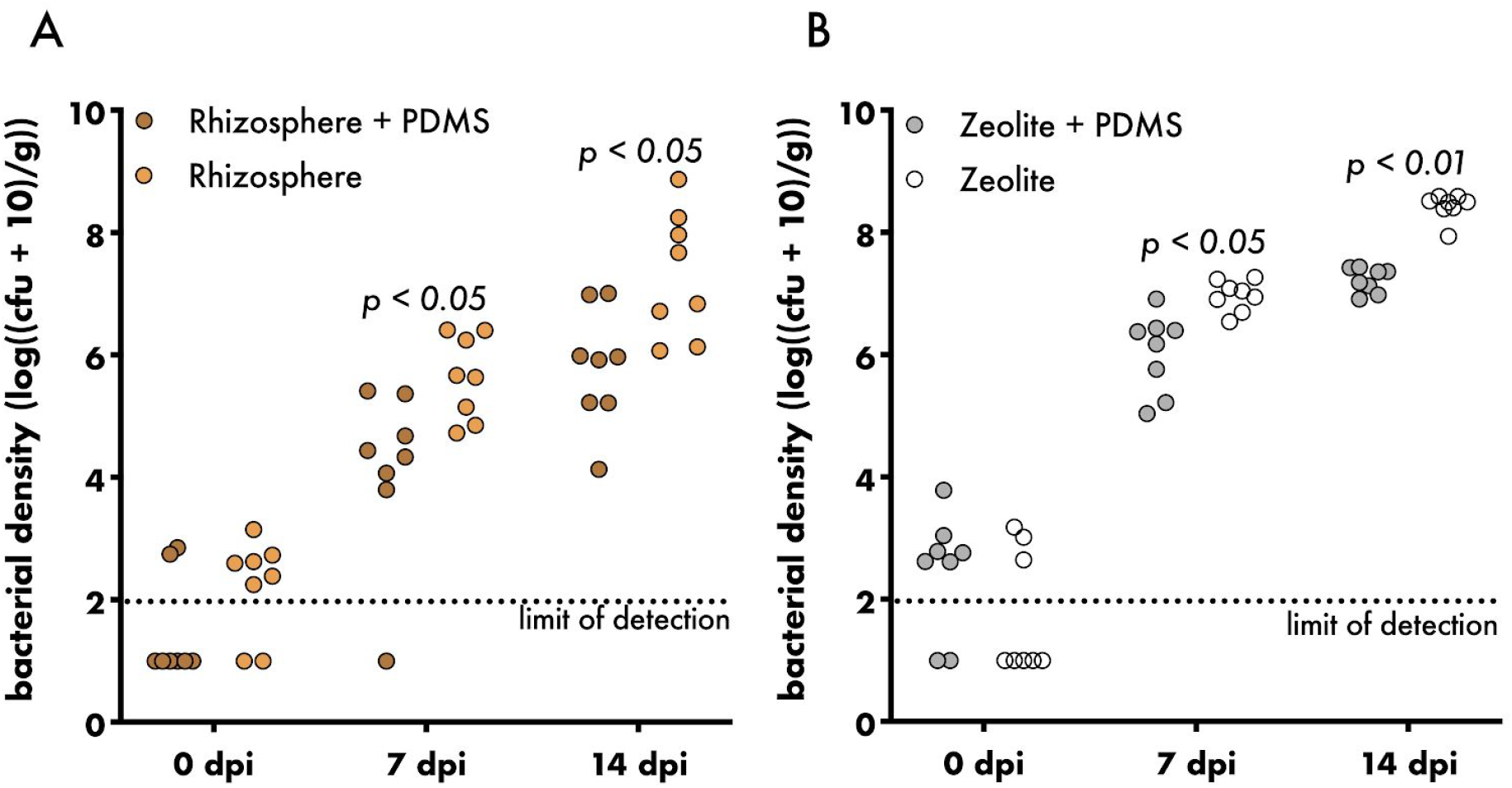
The effect of PDMS on bacterial load in the rhizosphere and zeolite. Bacterial density in the rhizosphere (**A**) or zeolite (**B**) of four-week-old plants inoculated with *Sphingomonas melonis* Fr1. Filled circles represent individual samples, dotted line represents limit of detection, Sidak’s multiple comparison test.

The Litterbox system allowed us to use a variety of molecular tools to investigate the intimate cross-talk between microbiota members and their host. To determine activation and shaping of the plant immune network, the RNA of the plant host was extracted, and the relative levels of plant immunity-associated hormones, salicylic acid (SA), ethylene (ET) and jasmonic acid (JA), were estimated using transcript markers, previously shown to reflect their respective hormone levels (Figure 5A-C) [59]. The presence of *P. syringae* DC3000 led to a stronger activation of the ethylene marker compared to *S. melonis* Fr1, even though *P. syringae* DC3000 was present in lower densities, and was previously shown to dampen ethylene production via its HopAF1 effector (Figure 5A,D) [72]. The reduction of the SA marker, pathogenesis-related 1 (PR1), within the first two days of *P. syringae* DC3000 inoculation, as well as the induction of PR1 following *S. melonis* Fr1 inoculation, has been shown previously (Figure 5B) [34]. Transcript levels of the JA marker were downregulated three hours after inoculation with *P. syringae* DC3000 and *S. melonis* Fr1, but upregulated at 48 hours after inoculation (Figure 5C), highlighting the need to study plant responses in a time-dependent manner.

**Figure 5:**
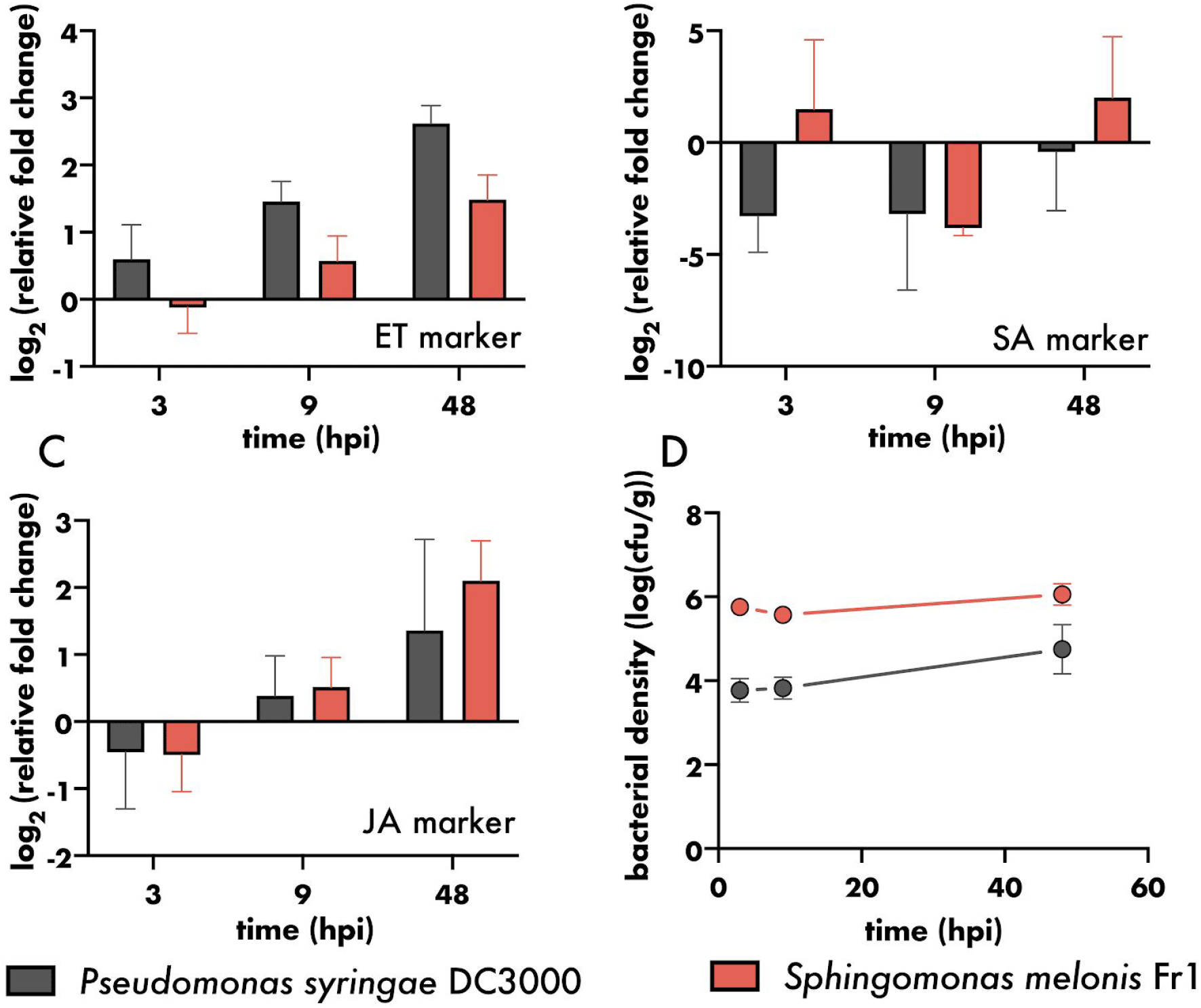
Plant responses to different bacteria. **A-C.** Log_2_ fold change in gene expression relative to mock treated control of (**A**) ethylene marker gene (At2g41230), (**B**) salicylic acid marker gene (PR1) and (**C**) jasmonic acid marker gene (At3g50280). Bars represent the mean of three biological replicates, error bars depict standard deviation. Each biological replicate comprises eight leaves from two plants. **D.** Bacterial density on aboveground plant parts of six-week-old plants relative to plant weight (fresh weight of aboveground plant parts). Filled circles represent mean of six plants, error bars represent standard deviation.

The controlled growth conditions for Arabidopsis and associated plant-adapted microbes that the Litterbox system offers also allowed the study of microbe-microbe interactions at the single-cell resolution. For example, we observed the distribution of cell aggregates of *P. syringae* DC3000 and *S. melonis* Fr1 14 days after co-inoculation onto Arabidopsis leaves (Figure 6). Bacterial community members from leaves of environmentally grown Arabidopsis have been shown to establish non-random spatial patterns [60]. As the Litterbox system produces plants that resemble environmentally grown plants with comparable bacterial load, it is the ideal system to test the ecological concepts underlying these non-random spatial patterns under laboratory conditions [73]. Bioreporters and single-cell approaches are useful tools to broaden our understanding of bacterial adaptations, colonisation patterns and physiology in the phyllosphere [1,74–76]. Having a growth system that resembles natural conditions opens the opportunity to study bacterial adaptations to varying environmental factors in a controlled manner (*e.g.*, humidity, light intensity and exposure, and temperature).

**Figure 6:**
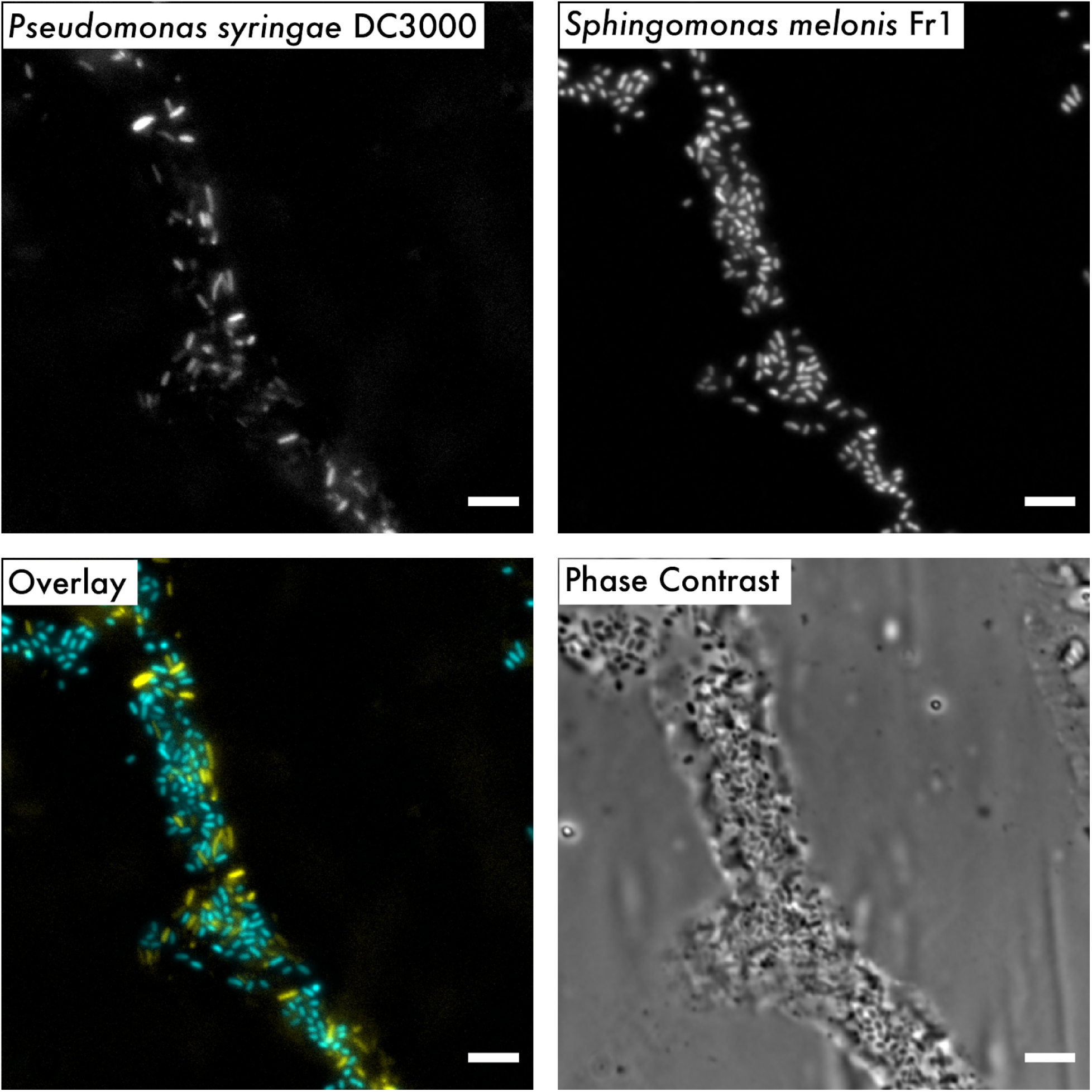
Bacterial distribution on leaves. Representative images of bacterial distribution on leaves after recovery using cuticle tape lifts. Fluorescently tagged *P. syringae* DC3000 and *S. meloni*s Fr1 were co-inoculated onto seven-week-old Arabidopsis leaves and sampled at 14 days post-inoculation. Overlay image represents the combined pseudo-coloured micrographs of each strain (yellow: *P. syringae* DC3000, cyan: *S. melonis* Fr1). Bar scale 5 µm.

## Conclusion

Zeolite can be easily sourced from readily available cat litter making the Litterbox system affordable and easy to establish. The low variation and consistency in plant growth allows for predictable and reproducible experiments, alleviating the chance of false positive and false negative measurements. The Litterbox system is compact, thereby enabling experiments with large sample sizes. Most importantly, plant growth and leaf bacterial densities resemble those found in temperate environments, therefore enabling the plant microbiota research community to investigate plant-microbe interactions and microbe-microbe interactions, and their underlying mechanisms, in an environmental context, while being able to control environmental factors (*e.g.*, humidity, light intensity and exposure, temperature), microbial community composition and plant genotypes. We anticipate that this versatile and affordable gnotobiotic plant growth system will advance microbiota research and broaden our understanding of microbiota assembly, composition and its effects on the plant host.

## Acknowledgments

The authors thank Michal Bernach for sharing his expertise about PDMS and Alan Woods for his technical support.

## Author Contributions

Conceptualization, M.M. and M.R-E.; methodology, M.M., R.S., J.C., P.J., M.R-E.; formal analysis, M.M.; investigation, M.M., R.S.; writing—original draft preparation, M.M.; writing—review and editing, M.R-E., R.S., P.J., J.C.; visualization, M.M., R.S.; supervision, M.R-E, P.J.; project administration, M.R-E.; funding acquisition, M.R-E, P.J.. All authors have read and agreed to the published version of the manuscript.

## References

1. Remus-Emsermann, M.N.P.; Schlechter, R.O. Phyllosphere microbiology: at the interface between microbial individuals and the plant host. New Phytol. 2018, 218, 1327–1333.

2. Schlechter, R.O.; Miebach, M.; Remus-Emsermann, M.N.P. Driving factors of epiphytic bacterial communities: A review. J. Adv. Res. 2019, 19, 57–65.

3. Bulgarelli, D.; Schlaeppi, K.; Spaepen, S.; Ver Loren van Themaat, E.; Schulze-Lefert, P. Structure and functions of the bacterial microbiota of plants. Annu. Rev. Plant Biol. 2013, 64, 807–838.

4. Vandenkoornhuyse, P.; Quaiser, A.; Duhamel, M.; Le Van, A.; Dufresne, A. The importance of the microbiome of the plant holobiont. New Phytol. 2015, 206, 1196–1206.

5. Sánchez-Cañizares, C.; Jorrín, B.; Poole, P.S.; Tkacz, A. Understanding the holobiont: the interdependence of plants and their microbiome. Curr. Opin. Microbiol. 2017, 38, 188–196.

6. Hassani, M.A.; Durán, P.; Hacquard, S. Microbial interactions within the plant holobiont. Microbiome 2018, 6, 58.

7. Chen, M.; Arato, M.; Borghi, L.; Nouri, E.; Reinhardt, D. Beneficial services of arbuscular mycorrhizal fungi - from ecology to application. Front. Plant Sci. 2018, 9.

8. Wang, Q.; Liu, J.; Zhu, H. Genetic and molecular mechanisms underlying symbiotic specificity in legume-*Rhizobium* interactions. Front. Plant Sci. 2018, 9, 313.

9. Innerebner, G.; Knief, C.; Vorholt, J.A. Protection of *Arabidobsis thaliana* against leaf-pathogenic *Pseudomonas syringae* by *Sphingomonas strains* in a controlled model system. Appl. Environ. Microbiol. 2011, 77, 3202–3210.

10. Ritpitakphong, U.; Falquet, L.; Vimoltust, A.; Berger, A.; Métraux, J.-P.; L’Haridon, F. The microbiome of the leaf surface of Arabidopsis protects against a fungal pathogen. New Phytol. 2016, 210, 1033–1043.

11. Berg, M.; Koskella, B. Nutrient- and dose-dependent microbiome-mediated protection against a plant pathogen. Curr. Biol. 2018, 28, 2487–2492.e3.

12. Spaepen, S.; Vanderleyden, J.; Okon, Y. Chapter 7 Plant growth-promoting actions of Rhizobacteria. In Adv. Bot. Res.; Academic Press, 2009; Vol. 51, pp. 283–320.

13. Lau, J.A.; Lennon, J.T. Rapid responses of soil microorganisms improve plant fitness in novel environments. Proc. Natl. Acad. Sci. U. S. A. 2012, 109, 14058–14062.

14. Zengerer, V.; Schmid, M.; Bieri, M.; Müller, D.C.; Remus-Emsermann, M.N.P.; Ahrens, C.H.; Pelludat, C. *Pseudomonas orientalis* F9: a potent antagonist against phytopathogens with phytotoxic effect in the apple flower. Front. Microbiol. 2018, 9, 145.

15. Kong, Z.; Hart, M.; Liu, H. Paving the way from the lab to the field: using synthetic microbial consortia to produce high-quality crops. Front. Plant Sci. 2018, 9, 1467.

16. Sergaki, C.; Lagunas, B.; Lidbury, I.; Gifford, M.L.; Schäfer, P. Challenges and approaches in microbiome research: from fundamental to applied. Front. Plant Sci. 2018, 9, 1205.

17. Busby, P.E.; Soman, C.; Wagner, M.R.; Friesen, M.L.; Kremer, J.; Bennett, A.; Morsy, M.; Eisen, J.A.; Leach, J.E.; Dangl, J.L. Research priorities for harnessing plant microbiomes in sustainable agriculture. PLoS Biol. 2017, 15, e2001793.

18. Delmotte, N.; Knief, C.; Chaffron, S.; Innerebner, G.; Roschitzki, B.; Schlapbach, R.; von Mering, C.; Vorholt, J.A. Community proteogenomics reveals insights into the physiology of phyllosphere bacteria. Proc. Natl. Acad. Sci. U. S. A. 2009, 106, 16428–16433.

19. Knief, C.; Delmotte, N.; Chaffron, S.; Stark, M.; Innerebner, G.; Wassmann, R.; von Mering, C.; Vorholt, J.A. Metaproteogenomic analysis of microbial communities in the phyllosphere and rhizosphere of rice. ISME J. 2012, 6, 1378–1390.

20. Levy, A.; Salas Gonzalez, I.; Mittelviefhaus, M.; Clingenpeel, S.; Herrera Paredes, S.; Miao, J.; Wang, K.; Devescovi, G.; Stillman, K.; Monteiro, F.; et al. Genomic features of bacterial adaptation to plants. Nat. Genet. 2018, 50, 138–150.

21. Turner, T.R.; Ramakrishnan, K.; Walshaw, J.; Heavens, D.; Alston, M.; Swarbreck, D.; Osbourn, A.; Grant, A.; Poole, P.S. Comparative metatranscriptomics reveals kingdom level changes in the rhizosphere microbiome of plants. ISME J. 2013, 7, 2248–2258.

22. Makiola, A.; Dickie, I.A.; Holdaway, R.J.; Wood, J.R.; Orwin, K.H.; Glare, T.R. Land use is a determinant of plant pathogen alpha-but not beta-diversity. Mol. Ecol. 2019, 28, 3786–3798.

23. Vorholt, J.A.; Vogel, C.; Carlström, C.I.; Müller, D.B. Establishing causality: opportunities of synthetic communities for plant microbiome research. Cell Host Microbe 2017, 22, 142–155.

24. Bai, Y.; Müller, D.B.; Srinivas, G.; Garrido-Oter, R.; Potthoff, E.; Rott, M.; Dombrowski, N.; Münch, P.C.; Spaepen, S.; Remus-Emsermann, M.; et al. Functional overlap of the Arabidopsis leaf and root microbiota. Nature 2015, 528, 364–369.

25. Liu, Y.-X.; Qin, Y.; Bai, Y. Reductionist synthetic community approaches in root microbiome research. Curr. Opin. Microbiol. 2019, 49, 97–102.

26. Carlström, C.I.; Field, C.M.; Bortfeld-Miller, M.; Müller, B.; Sunagawa, S.; Vorholt, J.A. Synthetic microbiota reveal priority effects and keystone strains in the Arabidopsis phyllosphere. Nat Ecol Evol 2019, 3, 1445–1454.

27. Stockwell, V.O.; Johnson, K.B.; Sugar, D.; Loper, J.E. Control of fire blight by *Pseudomonas fluorescens* A506 and *Pantoea vagans* C9-1 applied as single strains and mixed inocula. Phytopathology 2010, 100, 1330–1339.

28. Stockwell, V.O.; Johnson, K.B.; Sugar, D.; Loper, J.E. Mechanistically compatible mixtures of bacterial antagonists improve biological control of fire blight of pear. Phytopathology 2011, 101, 113–123.

29. Berens, M.L.; Wolinska, K.W.; Spaepen, S.; Ziegler, J.; Nobori, T.; Nair, A.; Krüler, V.; Winkelmüller, T.M.; Wang, Y.; Mine, A.; et al. Balancing trade-offs between biotic and abiotic stress responses through leaf age-dependent variation in stress hormone cross-talk. Proc. Natl. Acad. Sci. U. S. A. 2019, 116, 2364–2373.

30. Finkel, O.M.; Salas-González, I.; Castrillo, G.; Spaepen, S.; Law, T.F.; Teixeira, P.J.P.L.; Jones, C.D.; Dangl, J.L. The effects of soil phosphorus content on plant microbiota are driven by the plant phosphate starvation response. PLoS Biol. 2019, 17, e3000534.

31. Fierer, N.; Jackson, R.B. The diversity and biogeography of soil bacterial communities. Proc. Natl. Acad. Sci. U. S. A. 2006, 103, 626–631.

32. Andrew, D.R.; Fitak, R.R.; Munguia-Vega, A.; Racolta, A.; Martinson, V.G.; Dontsova, K. Abiotic factors shape microbial diversity in Sonoran Desert soils. Appl. Environ. Microbiol. 2012, 78, 7527–7537.

33. Tao, Y.; Xie, Z.; Chen, W.; Glazebrook, J.; Chang, H.-S.; Han, B.; Zhu, T.; Zou, G.; Katagiri, F. Quantitative nature of Arabidopsis responses during compatible and incompatible interactions with the bacterial pathogen *Pseudomonas syringae*. Plant Cell 2003, 15, 317–330.

34. Vogel, C.; Bodenhausen, N.; Gruissem, W.; Vorholt, J.A. The Arabidopsis leaf transcriptome reveals distinct but also overlapping responses to colonization by phyllosphere commensals and pathogen infection with impact on plant health. New Phytol. 2016, 212, 192–207.

35. Vogel, C.; Innerebner, G.; Zingg, J.; Guder, J.; Vorholt, J.A. Forward genetic *in planta screen for identification of plant-protective traits of Sphingomonas* sp. strain Fr1 against *Pseudomonas syringae* DC3000. Appl. Environ. Microbiol. 2012, 78, 5529–5535.

36. Henry, A.; Doucette, W.; Norton, J.; Jones, S.; Chard, J.; Bugbee, B. An axenic plant culture system for optimal growth in long-term studies. J. Environ. Qual. 2006, 35, 590–598.

37. Gunning, T.; Cahill, D.M. A soil-free plant growth system to facilitate analysis of plant pathogen interactions in roots. J. Phytopathol. 2009, 157, 497–501.

38. Koch, K.; Hartmann, K.D.; Schreiber, L.; Barthlott, W.; Neinhuis, C. Influences of air humidity during the cultivation of plants on wax chemical composition, morphology and leaf surface wettability. Environ. Exp. Bot. 2006, 56, 1–9.

39. Kosma, D.K.; Bourdenx, B.; Bernard, A.; Parsons, E.P.; Lü, S.; Joubès, J.; Jenks, M.A. The impact of water deficiency on leaf cuticle lipids of Arabidopsis. Plant Physiol. 2009, 151, 1918–1929.

40. Reisberg, E.E.; Hildebrandt, U.; Riederer, M.; Hentschel, U. Distinct phyllosphere bacterial communities on Arabidopsis wax mutant leaves. PLoS One 2013, 8, e78613.

41. Beattie, G.A. Water relations in the interaction of foliar bacterial pathogens with plants. Annu. Rev. Phytopathol. 2011, 49, 533–555.

42. Shaw, L.J.; Beaton, Y.; Glover, L.A.; Killham, K.; Meharg, A.A. Re-inoculation of autoclaved soil as a non-sterile treatment for xenobiotic sorption and biodegradation studies. Appl. Soil Ecol. 1999, 11, 217–226.

43. Bank, T.L.; Kukkadapu, R.K.; Madden, A.S.; Ginder-Vogel, M.A.; Baldwin, M.E.; Jardine, P.M. Effects of gamma-sterilization on the physico-chemical properties of natural sediments. Chem. Geol. 2008, 251, 1–7.

44. Lebeis, S.L.; Paredes, S.H.; Lundberg, D.S.; Breakfield, N.; Gehring, J.; McDonald, M.; Malfatti, S.; Glavina del Rio, T.; Jones, C.D.; Tringe, S.G.; et al. Salicylic acid modulates colonization of the root microbiome by specific bacterial taxa. Science 2015, 349, 860–864.

45. Zhang, J.; Liu, Y.-X.; Zhang, N.; Hu, B.; Jin, T.; Xu, H.; Qin, Y.; Yan, P.; Zhang, X.; Guo, X.; et al. NRT1.1B is associated with root microbiota composition and nitrogen use in field-grown rice. Nat. Biotechnol. 2019, 37, 676–684.

46. Adams, C.; Jacobson, A.; Bugbee, B. Ceramic aggregate sorption and desorption chemistry: implications for use as a component of soilless media. J. Plant Nutr. 2014, 37, 1345–1357.

47. Ming, D.W.; Allen, E.R. Use of natural zeolites in agronomy, horticulture and environmental soil remediation. Rev. Mineral. Geochem. 2001, 45, 619–654.

48. Iskander, A.L.; Khald, E.M.; Sheta, A.S. Zinc and manganese sorption behavior by natural zeolite and bentonite. Sci. Ann. Univ. Agric. Sci. Vet. Med. 2011, 56, 43–48.

49. Research and Application | ZeoponiX Inc Available online: http://zeoponix.com/?page_id=94 (accessed on Jan 8, 2020).

50. Lundberg, D.S.; Lebeis, S.L.; Paredes, S.H.; Yourstone, S.; Gehring, J.; Malfatti, S.; Tremblay, J.; Engelbrektson, A.; Kunin, V.; Del Rio, T.G.; et al. Defining the core *Arabidopsis thaliana* root microbiome. Nature 2012, 488, 86–90.

51. Peyraud, R.; Kiefer, P.; Christen, P.; Massou, S.; Portais, J.-C.; Vorholt, J.A. Demonstration of the ethylmalonyl-CoA pathway by using 13C metabolomics. Proc. Natl. Acad. Sci. U. S. A. 2009, 106, 4846–4851.

52. Jameson, P.E.; Morris, R.O. Zeatin-like cytokinins in yeast: detection by immunological methods. J. Plant Physiol. 1989, 135, 385–390.

53. Schlechter, R.O.; Jun, H.; Bernach, M.; Oso, S.; Boyd, E.; Muñoz-Lintz, D.A.; Dobson, R.C.J.; Remus, D.M.; Remus-Emsermann, M.N.P. Chromatic Bacteria - a broad host-range plasmid and chromosomal insertion toolbox for fluorescent protein expression in bacteria. Front. Microbiol. 2018, 9, 3052.

54. Schlechter, R.; Remus-Emsermann, M. Delivering “Chromatic Bacteria” fluorescent protein tags to Proteobacteria using conjugation. Bio Protoc. 2019, 9.

55. Buonaurio, R.; Stravato, V.M.; Kosako, Y.; Fujiwara, N.; Naka, T.; Kobayashi, K.; Cappelli, C.; Yabuuchi, E. *Sphingomonas melonis* sp. nov., a novel pathogen that causes brown spots on yellow Spanish melon fruits. Int. J. Syst. Evol. Microbiol. 2002, 52, 2081–2087.

56. Cuppels, D.A. Generation and characterization of Tn5 insertion mutations in *Pseudomonas syringae* pv. tomato. Appl. Environ. Microbiol. 1986, 51, 323–327.

57. Nolan, T.; Huggett, J.F.; Sanchez, E. Good practice guide for the application of quantitative PCR (qPCR). LGC 2013.

58. Czechowski, T.; Stitt, M.; Altmann, T.; Udvardi, M.K.; Scheible, W.-R. Genome-wide identification and testing of superior reference genes for transcript normalization in Arabidopsis. Plant Physiol. 2005, 139, 5–17.

59. Kim, Y.; Tsuda, K.; Igarashi, D.; Hillmer, R.A.; Sakakibara, H.; Myers, C.L.; Katagiri, F. Mechanisms underlying robustness and tunability in a plant immune signaling network. Cell Host Microbe 2014, 15, 84–94.

60. Remus-Emsermann, M.N.P.; Lücker, S.; Müller, D.B.; Potthoff, E.; Daims, H.; Vorholt, J.A. Spatial distribution analyses of natural phyllosphere-colonizing bacteria on *Arabidopsis thaliana* revealed by fluorescence *in situ* hybridization. Environ. Microbiol. 2014, 16, 2329–2340.

61. Schindelin, J.; Arganda-Carreras, I.; Frise, E.; Kaynig, V.; Longair, M.; Pietzsch, T.; Preibisch, S.; Rueden, C.; Saalfeld, S.; Schmid, B.; et al. Fiji: an open-source platform for biological-image analysis. Nat. Methods 2012, 9, 676–682.

62. Roué, J.; Chauvet, H.; Brunel-Michac, N.; Bizet, F.; Moulia, B.; Badel, E.; Legué, V. Root cap size and shape influence responses to the physical strength of the growth medium in *Arabidopsis thaliana* primary roots. J. Exp. Bot. 2019.

63. Eapen, D.; Barroso, M.L.; Ponce, G.; Campos, M.E.; Cassab, G.I. Hydrotropism: root growth responses to water. Trends Plant Sci. 2005, 10, 44–50.

64. Remus-Emsermann, M.N.P.; Pelludat, C.; Gisler, P. Conjugation dynamics of self-transmissible and mobilisable plasmids into *E. coli* O157: H7 on *Arabidopsis thaliana* rosettes. bioRxiv 2018.

65. Vogel, C.; Innerebner, G.; Zingg, J.; Guder, J.; Vorholt, J.A. Forward genetic *in planta screen for identification of plant-protective traits of Sphingomonas* sp. strain Fr1 against *Pseudomonas syringae* DC3000. Appl. Environ. Microbiol. 2012, 78, 5529–5535.

66. Burch, A.Y.; Do, P.T.; Sbodio, A.; Suslow, T.V.; Lindow, S.E. High-level culturability of epiphytic bacteria and frequency of biosurfactant producers on leaves. Appl. Environ. Microbiol. 2016, 82, 5997–6009.

67. Gekenidis, M.-T.; Gossin, D.; Schmelcher, M.; Schöner, U.; Remus-Emsermann, M.N.P.; Drissner, D. Dynamics of culturable mesophilic bacterial communities of three fresh herbs and their production environment. J. Appl. Microbiol. 2017, 123, 916–932.

68. Kremer, J.M.; Paasch, B.C.; Rhodes, D.; Thireault, C.; Froehlich, J.E.; Schulze-Lefert, P.; Tiedje, J.M.; He, S.Y. FlowPot axenic plant growth system for microbiota research. bioRxiv 2018, 254953.

69. Van der Ent, S.; Van Hulten, M.; Pozo, M.J.; Czechowski, T.; Udvardi, M.K.; Pieterse, C.M.J.; Ton, J. Priming of plant innate immunity by rhizobacteria and β-aminobutyric acid: differences and similarities in regulation. New Phytol. 2009, 183, 419–431.

70. Pieterse, C.M.J.; Zamioudis, C.; Berendsen, R.L.; Weller, D.M.; Van Wees, S.C.M.; Bakker, P.A.H.M. Induced systemic resistance by beneficial microbes. Annu. Rev. Phytopathol. 2014, 52, 347–375.

71. Raaijmakers, J.M.; Leeman, M.; Van Oorschot, M.M.P.; Van der Sluis, I.; Schippers, B.; Bakker, P. Dose-response relationships in biological control of *Fusarium* wilt of radish by *Pseudomonas* spp. Phytopathology 1995, 85, 1075–1080.

72. Washington, E.J.; Mukhtar, M.S.; Finkel, O.M.; Wan, L.; Banfield, M.J.; Kieber, J.J.; Dangl, J.L. *Pseudomonas syringae* type III effector HopAF1 suppresses plant immunity by targeting methionine recycling to block ethylene induction. Proc. Natl. Acad. Sci. U. S. A. 2016, 113, E3577–86.

73. Meyer, K.M.; Leveau, J.H.J. Microbiology of the phyllosphere: a playground for testing ecological concepts. Oecologia 2012, 168, 621–629.

74. Leveau, J.H.; Lindow, S.E. Appetite of an epiphyte: quantitative monitoring of bacterial sugar consumption in the phyllosphere. Proc. Natl. Acad. Sci. U. S. A. 2001, 98, 3446–3453.

75. Monier, J.-M.; Lindow, S.E. Frequency, size, and localization of bacterial aggregates on bean leaf surfaces. Appl. Environ. Microbiol. 2004, 70, 346–355.

76. Sandhu, A.; Halverson, L.J.; Beattie, G.A. Bacterial degradation of airborne phenol in the phyllosphere. Environ. Microbiol. 2007, 9, 383–392.

