## Supplementary figures for "Litterbox - A gnotobiotic zeolite-clay system to investigate Arabidopsis-microbe interactions"

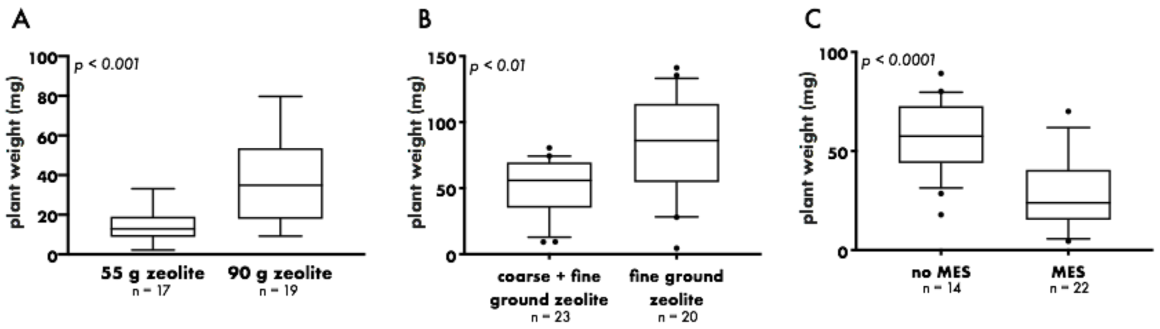


**Figure S1: Optimisation of the Litterbox system. A.** Effect of zeolite amount on plant weight (fresh weight aboveground plant parts) of six-week-old plants grown on coarse (35 g or 70 g) zeolite covered by fine ground (20 g) zeolite. **B.** Effect of zeolite granularity on plant weight of six-week-old plants grown on 90 g of zeolite (70 + 20 g and 90 g). **C.** Effect of MES buffer (2.5 mM) on plant weight of six-week-old plants grown on coarse zeolite (70 g) covered by fine ground zeolite (20 g). Tukey's boxplots.

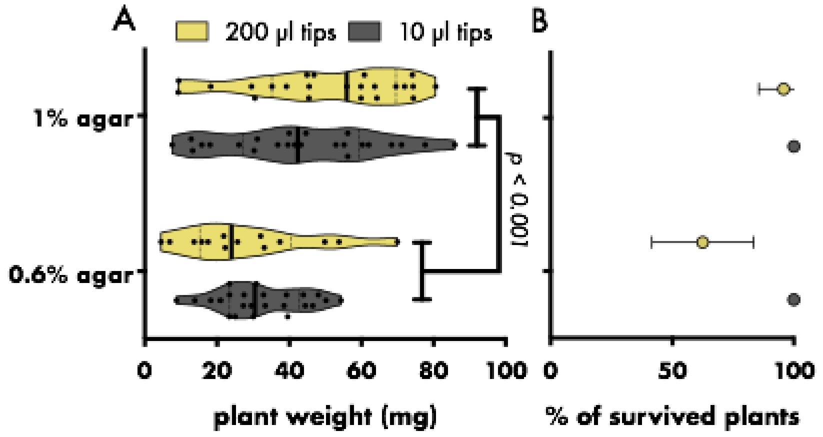


**Figure S2: Effect of agar concentration and tip size of agar filled pipette tips on plant growth. A.** Plant weight of six-week-old plants (fresh weight aboveground plant parts) of plants sown either on cut 10 µl or 200 µl pipette tips filled with either 0.6% or 1% agar. Seedlings were transferred seven days after sowing to 70 g coarse zeolite covered by 20 g fine ground zeolite. **B.** Percentage of plants grown to full size per transplanted seedling. Filled circles represent sample mean, error bars depict standard deviation, dots mark individual plants, thick bar represents median, dotted bars represent quartiles, two-way ANOVA.

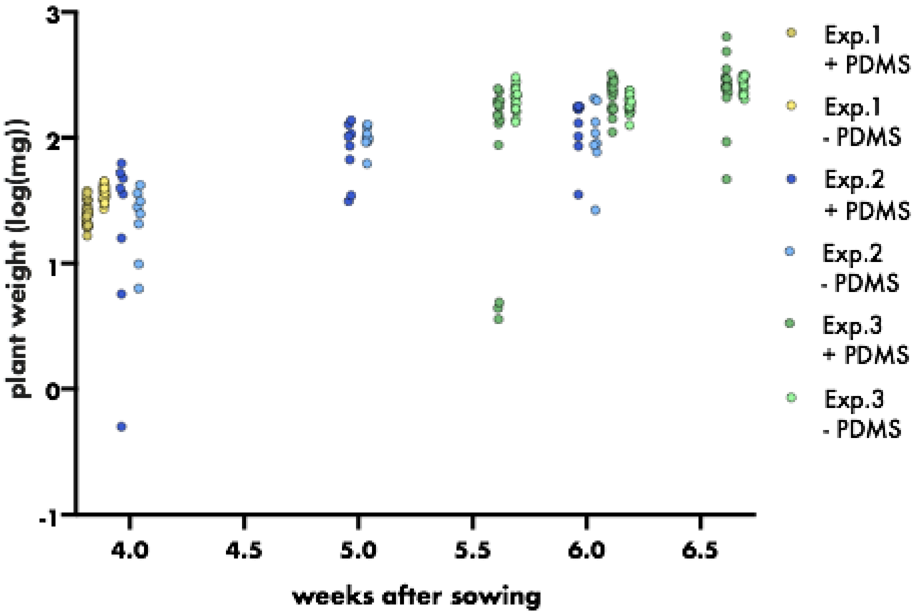


**Figure S3: Effect of PDMS sheet on plant weight.** Plant weight (fresh weight aboveground plant parts) at varying ages, grown either with or without PDMS covering the substrate. Plants were grown on 90 g finely ground zeolite. Dots represent individual plants, different colors mark independent experiments, dark color = + PDMS, light color = - PDMS.
